# A rapid agarose-free protocol for preparing human organotypic lung cultures to study respiratory virus infection and evaluate antivirals *ex vivo*

**DOI:** 10.64898/2026.02.12.705542

**Authors:** Lola Canus, Florentine Jacolin, Virginie Vasseur, Adeline Cezard, Eva Ogire, Anne Aublin-Gex, Amelie Bourgeais, Camille David, Alexandra Erny, Fabienne Archer, Antoine Legras, Damien Sizaret, Antoine Guillon, Vincent Lotteau, Pierre-Olivier Vidalain, Mustapha Si-Tahar, Laure Perrin-Cocon, Cyrille Mathieu

## Abstract

We describe a standardized and reproducible procedure to generate human organotypic lung cultures from surgical lung resection for the study of respiratory infections. The protocol details tissue harvesting, biopsy punching, mechanical slicing, culture at the air–liquid interface. This technique enables robust *ex vivo* infections of human lung tissue with respiratory viruses, including Influenza A and Nipah. The described system can be used to study host-pathogen interactions, analyze innate immune responses, and evaluate antiviral candidates in physiologically relevant human lung tissue.

For complete details on the use and execution of this protocol, please refer to Cezard et al^1^.

## Before you begin

According to the World Health Organization, airborne viral infections are among the leading causes of human disease worldwide, with lower respiratory tract infections ranking among the deadliest. Understanding respiratory virus infection and host response at the tissue level remains challenging due to the limitations of experimental models. *In vivo* animal models, such as mice and hamsters, are widely used but often remain poorly suited for tissue-specific mechanistic investigations and are constrained by experimental complexity and ethical considerations. In contrast, *in vitro* systems enable controlled experimentation but fail to recapitulate the three-dimensional architecture and cellular diversity of the lung. Organotypic lung cultures therefore represent a powerful *ex vivo* alternative, as they preserve native tissue organization and, when maintained at the air–liquid interface, reproduce key physiological features of the respiratory epithelium. We have previously established standardized organotypic cultures from animals, including brain substructures, lung, kidney, and liver for infection studies. Our models enable reproducible *ex vivo* studies of zoonotic infections and provide insights into tissue-specific viral tropism and innate immune responses while contributing to the reduction of animal experimentation^1–9^. Building on this experience, we have implemented this approach using human lung tissue.

This protocol describes a rapid and standardized method to generate and maintain human organotypic lung cultures (huOLC) from surgical resections and to infect huOLC with various respiratory pathogens such as Influenza virus and Nipah virus. Compared with previously reported protocols, our method introduces and combines several key innovations designed to improve tissue survival, metabolic respiration, and the maintenance of resident immune cells for at least the duration of infection studies^10^. It details key steps from tissue coring and slicing, huOLC maintenance, and infection with BSL-2 and BSL-4 pathogens with the output tests detailed to follow in-depth examination of viral infection dynamics, host tissue response as well as the evaluation of antivirals. Please note that this protocol necessitates proper institutional and ethical approval prior to working with human tissue samples.

## Innovation

This workflow describes an agarose-free protocol to generate huOLCs from fresh surgical resections. Unlike many studies in which the lung tissue is inflated and embedded in low-melting agarose (1-3%) prior to slicing with vibratome or Krumdieck slicers^11–21^, our protocol enables immediate processing and thus significantly accelerates the experimental workflow. Tissue slicing is performed using a tissue chopper, preserving native tissue organization, while culture of huOLCs at the air-liquid interface on microporous and hydrophobic membranes^22^ better mimics physiological lung conditions *in vivo*. This configuration allows basolateral diffusion of nutrients while maintaining exposure of the apical surface to air. Unlike submerged culture conditions commonly used for organotypic lung slices^11,14–17,19–21,23^, this approach limits oxygen deprivation and reduces the loss of non-adherent cells, supporting sustained structural integrity and viability for at least 6 days of culture.

In previous studies, huOLCs have been maintained in various culture media. Dulbecco’s Modified Eagle Medium/F12 (DMEM/F12) ^12–15,24^ or HAM (DMEM/F12 HAM) ^16–20^ formulations generally support tissue viability and structural integrity, whereas Minimum Essential Medium (MEM) has been reported as less suitable for long-term maintenance of complex lung architecture^23^. In some studies, fetal bovine serum (FBS) is added at low concentrations (typically 0.1%) to help preserve cell composition from cell differentiation. Here, we have chosen a serum-free 1:1 mixture of Advanced DMEM/F-12 and Bronchial Epithelial Growth Medium (BEGM), the latter being specifically formulated for airway epithelial cells. This combination provides optimal epithelial preservation while maintaining low basal secretion of inflammatory interleukin (IL)-6 making it particularly suited for infection studies^25^.

## Institutional permissions

Human lung resections:

- Centre Hospitalier Régional Universitaire de Tours (CHRU): Informed consent was obtained from all patients in accordance with the Helsinki Declaration and the ethical principles set out in the Belmont Report of the U.S. Department of Health and Human Services.
- Tissue and cell collections were declared to the French Ministry of Higher Education, Research, and Innovation (MESRI, authorization DC-2008-308). All experimental procedures complied with the Code of Ethics of the World Medical Association and were approved by the Ethics Committee of the CHRU of Tours.
- Hospices Civils de Lyon (HCL): Informed consent was obtained from all patients in accordance with the Helsinki Declaration. Tissue and cell collections were declared to the French Ministry of Higher Education, Research, and Innovation (MESRI, authorization DC-2008-308). Human biological samples and associated data were obtained from the HCL Biobank (CRB-HCL BB-0033-00046).

## Preparation of the culture plates with inserts

### Timing: 1-3 about 10 min per 6-well plate under biosafety cabinet in the BSL-2

1. Remove the lid of a sterile 6-well plate and add 1 mL of huOLC culture medium (1:1 mixture of supplemented BEGM and Advanced medium) to each well.
2. Open the packaging of the PTFE membrane inserts under sterile conditions.
3. Using straight and sterile tweezers, gently place one insert into each well, ensuring it lands on the surface of the medium. This step helps reverse the hydrophobicity of the PTFE membrane, allowing proper contact with the medium. **Critical step (1 and 2):** This should be performed one day before tissue slicing. **Critical step (3):** Ensure no air bubbles are trapped between the PTFE membrane and the medium, as this can impair reversion of the hydrophobicity of the membrane and liquid absorption by cultures throughout micropores, *in fine* affecting tissue viability and nutrients diffusion. **Pause point:** Inserts can be prepared in advance (e.g., on Friday for an experiment on Monday). If prepared more than 24 h in advance, replace the medium to maintain optimal conditions.
4. Incubate the prepared 6-well plate overnight at 37 °C in a humidified incubator with 5% CO₂.

## Slicing preparation

### Timing: 1-9 about 30 min

1. Decontaminate the laboratory bench thoroughly using 70% ethanol (EtOH).
2. Cover the bench with aluminum foil and spray it with 70% EtOH. Allow the aluminum foil to dry. **Critical step:** Place the aluminum foil with the dull side facing down to ensure proper surface contact. **Note:** The bench can be prepared the day before. Use of aluminum foil is only required when preparation is not performed in a clean white room.
3. Sterilize all slicing and dissection equipment with 70% EtOH, *i.e.* tissue chopper, Leica microscope. All the equipment must be fully dry before use.
4. Prepare a container filled with 70% EtOH for decontamination of dissection material before use (*i.e.* razor blades, scissors, curved tweezers, straight tweezers). After decontamination, put dissection tools on a sterile petri dish to dry before use.
5. Prepare the Whatman paper (6 cm x 6 cm). **Critical step:** Whatman paper must be autoclaved before use. **Note:** Autoclaving can be performed in advance and stored in a sterile container until use.
6. Prepare the slicing medium by combining Hibernate A + 10,000 U/mL Penicillin/Streptomycin in a 50 mL sterile falcon tube.
7. Aliquot 5 mL of slicing medium in 35 mm sterile petri dishes and keep the lid on the dishes as long as possible to avoid contamination. **Critical step:** Store the remaining slicing medium at 4°C.
8. Prepare the transfer pipette by cutting the conical extremity of a 5 mL sterile plastic pipette, remove the filter without touching the pipette and insert the top part into the pipette bulb.
9. Set up the tissue chopper by adjusting the slicing parameters for the desired thickness (here 500 µm), blade force and speed. **Caution:** Razor and scalpel blades are extremely sharp. Handle with care and wear appropriate protective equipment.

## Key resources table

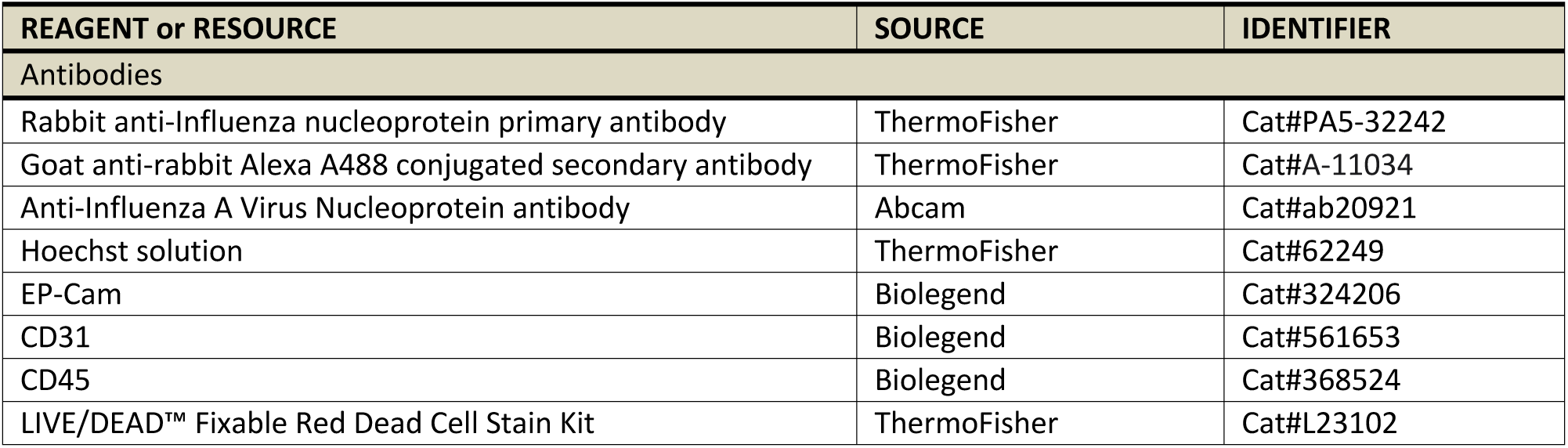
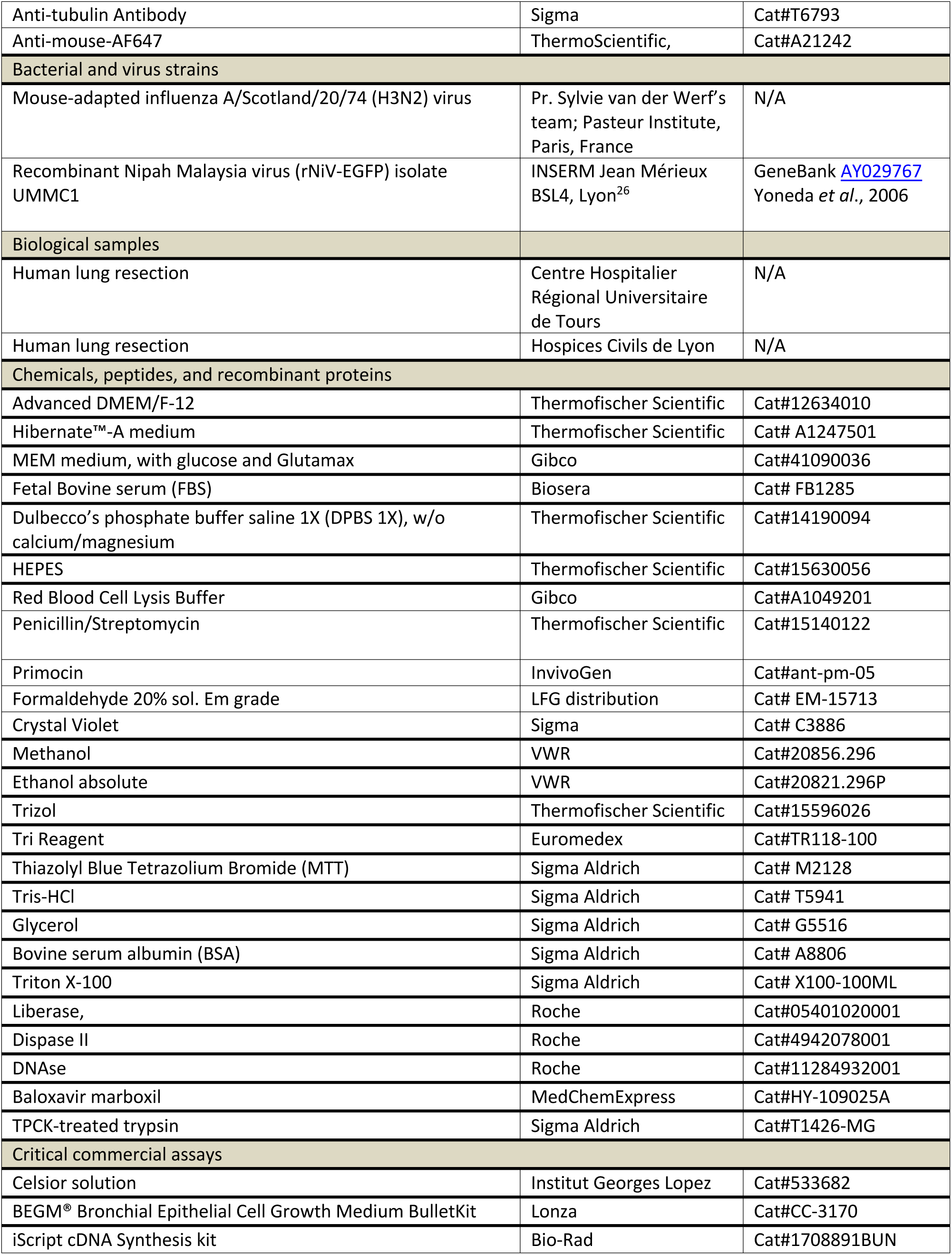
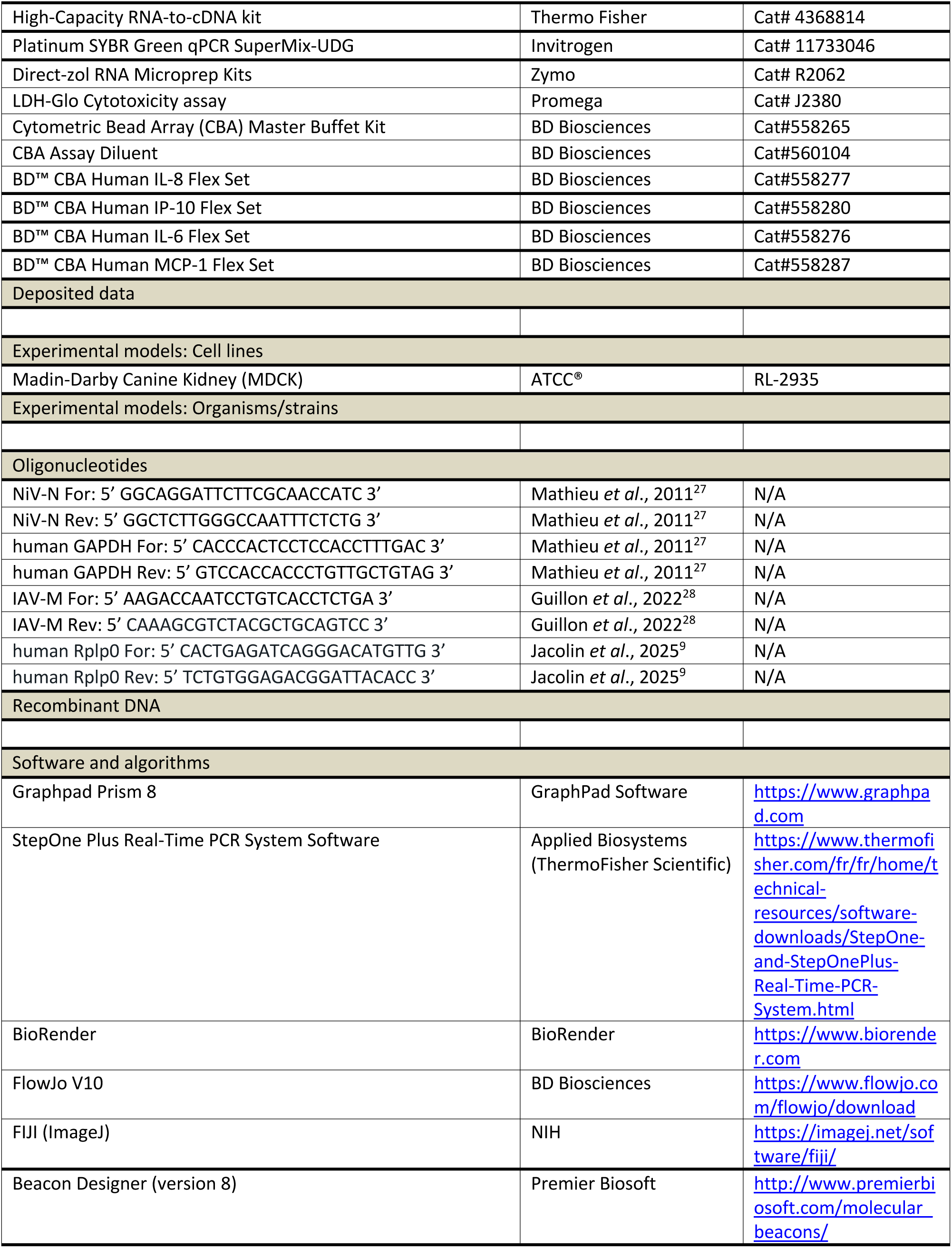
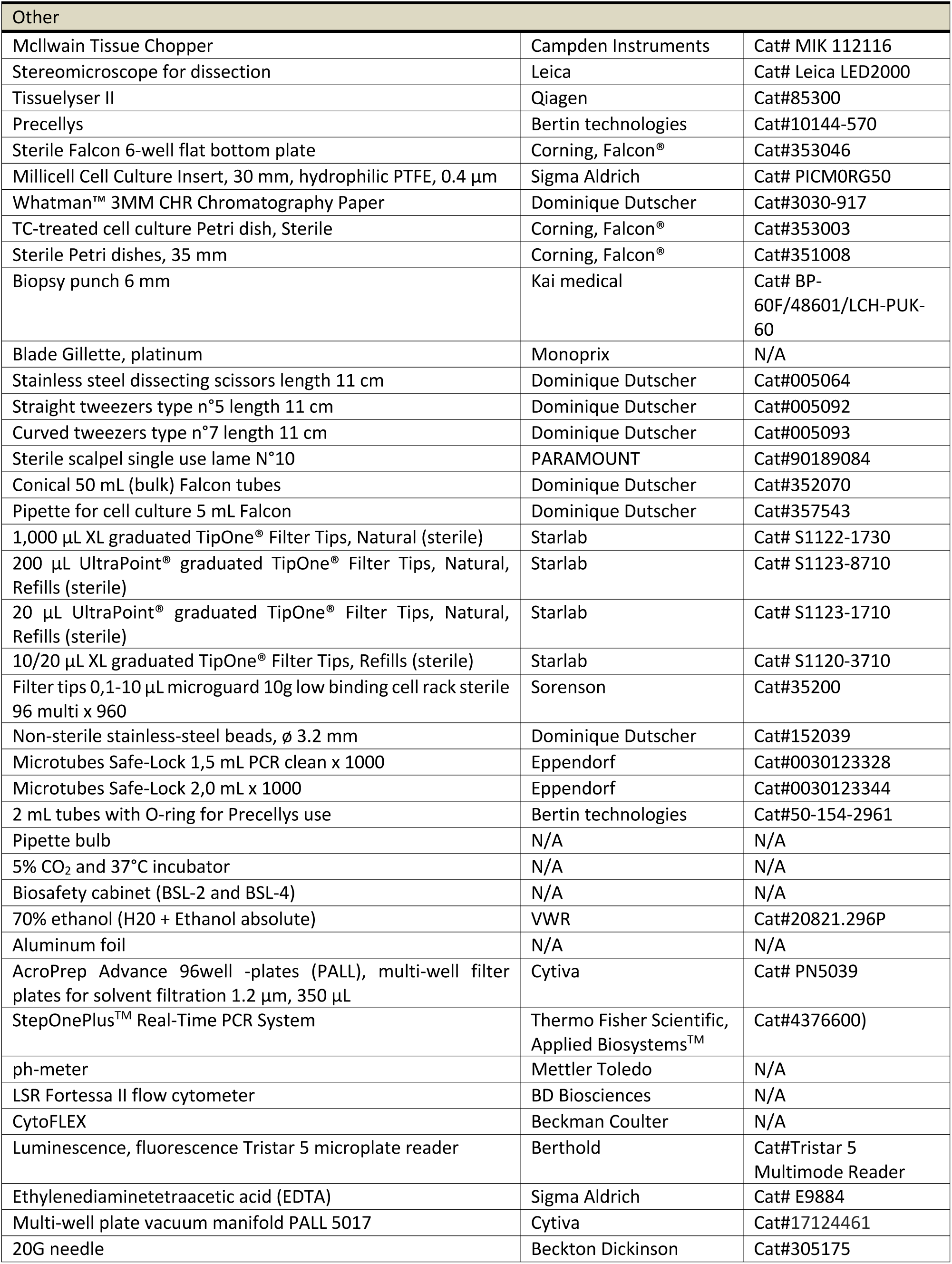

***N/A: Not applicable***

## Materials and equipment

### Supplemented BEGM

| Reagent | Final concentration | Amount |
| --- | --- | --- |
| BEBM Basal Medium | N/A | 500 mL |
| Bovine Pituitary Extract | 52 µg/mL | 2 mL |
| Insulin | 5 µg/mL | 500 µL |
| Hydrocortisone | 0.5 µg/mL | 500 µL |
| Gentamicin/ Amphotericin-1000 | Gentamicin: 30 µg/mL<br>Amphotericin: 15 ng/mL | 500 µL |
| Retinoic acid | 0.1 ng/mL | 500 µL |
| Transferrin | 10 µg/mL | 500 µL |
| Triiodothyronine | 6.5 ng/mL | 500 µL |
| Epinephrine | 0.5 µg/mL | 500 µL |
| Human Epidermal Growth Factor | 0.5 ng/mL | 500 µL |
| **Total** | **N/A** | **506 mL** |

N/A = not applicable.

**Note:** Divide the supplemented medium into 50 mL aliquots and store at –80°C for several months.

### Advanced DMEM/F12

Add 5 mL of Glutamax and 5 mL of HEPES to 500 ml of Advanced DMEM/F12. Store at 4°C up to 3 months.

### huOLC Culture medium

Mix 25 mL of BEGM supplemented with 25 mL of Advanced DMEM/F12 supplemented. Store up to 3 weeks at 4°C.

### MDCK (Madin-Darby Canine Kidney) complete culture medium

| Reagent | Final concentration | Amount |
| --- | --- | --- |
| MEM | N/A | 500 mL |
| Penicillin/ Streptomycin | 1% | 5 mL |
| FBS | 10% | 5 mL |
| **Total** | **N/A** | **510 mL** |

Store at 4°C up to 6 months.

### MDCK Infection medium

Add 1% of Penicillin/streptomycin 100X to for 500 mL of MEM medium. TPCK-treated trypsin is added extemporaneously with a final concentration of 1 µg/mL.

### Penicillin/streptomycin 100X

Once you receive the Penicillin/streptomycin make some aliquot and thaw the day you need it.

**Note:** Aliquot in 5 mL and store at –20°C until thawing process.

### Dissection/Slicing medium

Prepare 50 mL of Hibernate®-A supplemented with 500 µL of Penicillin/streptomycin 100X. Store at 4°C and only on the day of slicing (no long-term storage at 4°C or freezing possible).

### Formaldehyde 4%

Under a fume hood, dilute to 4% with DPBS 1X (10 mL of 20% formaldehyde with 40 mL of DPBS 1X). Store at –20°C until thawing process.

**Caution:** carcinogenic, mutagenic or toxic properties.

### Solution to saturate aldehyde groups after huOLC fixation

Weight 0.03 g glycine then add 30 mL DPBS 1X. Store at 4°C up to 1 year.

### Permeabilization solution for immunostaining

In 25 mL DPBS 1X, add 125 µL Triton X-100. Store at 4°C up to 1 year and mix before use.

### Blocking solution for immunostaining

| Reagent | Final concentration | Amount |
| --- | --- | --- |
| DPBS 1X | 1X | 20 mL |
| BSA | 1% | 0.2 g |
| Triton X-100 | 0.5% | 100 µL |
| **Total** | **N/A** | **20 mL** |

Store at 4°C up to 6 months and mix before use.

### LDH Storage Buffer

LDH storage buffer before is prepared according to [manufacturer instructions](https://www.promega.in/products/cell-health-assays/cell-viability-and-cytotoxicity-assays/ldh-glo-cytotoxicity-assay/?tabset0=1).

| Reagent | Final concentration | Amount |
| --- | --- | --- |
| Milli-Q water | N/A | 180 mL |
| Tris-HCl | 200 mM | 6.304 g |
| Glycerol | 10% | 20 mL |
| BSA | 1% | 2 g |
| **Total** | **N/A** | **200 mL** |

N/A = not applicable.

**Note:** Adjust pH after addition of Tris-HCl to pH 7.3. Store at at 4 °C, up to 6 months.

### Cytoflex digestion buffer

| Reagent | Final concentration | Amount |
| --- | --- | --- |
| DPBS 1X | N/A | 470 µL |
| Dispase II | 5 mg/mL | 2.5 mg |
| Liberase | 0.125 mg/mL | 25 µL |
| DNase | 0.1 mg/mL | 5 µL |
| **Total** | **N/A** | **500 µL** |

Store at 4°C up to 24 hours.

### Crystal Violet

| Reagent | Final concentration | Amount |
| --- | --- | --- |
| Crystal Violet | 0.3% | 3 g |
| Milli-Q water | N/A | 800 mL |
| Methanol | 80% | 200 mL |
| **Total** | **N/A** | **1 L** |

N/A = not applicable.

**Caution:** carcinogenic, mutagenic or toxic properties.

**Note:** Weigh crystal violet under a fume hood without aspiration, then transfer into a glass bottle. Turn on the fume hood aspiration. Carefully add Milli-Q water before adding methanol. Store at RT, away from light up to 1 year.

## Step-by-step method details

### Slicing procedure (Figure 1)

#### Timing: 1-3 about 10 min; 30 min for 4-5 (depending on the size of the resection) and finally 30 min for the last steps 6-10

In this step, we describe how to start the slicing procedure to ensure reproducible generation of uniform huOLC from lung resection, which is essential for preserving tissue viability and structural integrity for at least five days.

1. Collect the lung tissue samples immediately following surgical resection (performed at the CHRU of Tours or the HCL of Lyon) and transfer in Hibernate®-A containing primocin 0.1 mg/ml (or Penicillin/streptomycin) or in Celsior solution.
2. Transfer the lung resection into a large petri dish containing slicing medium. Gently rinse the resection to remove blood.
3. Transfer the lung resection to another petri dish containing slicing medium.
4. Carefully remove the pleura using a scalpel to facilitate slicing and trim the lung resection if necessary.
5. Use a sterile 6-mm biopsy punch fitted with a pipette bulb to generate cylindrical tissue cores from the lung resection. **Critical step:** This step should be performed by two people; one person stabilizes the lung resection using tweezers while the other one extracts the sample with the biopsy punch.
6. Place each “core tissue sample” in a 35-mm petri dish filled with a slicing medium. Let the core tissue sample gently elongate to ensure a proper form for slicing.
7. Once all the core tissue samples are prepared, transfer one tissue core onto 4 autoclaved Whatman papers with a drop of slicing medium.
8. Place the Whatman papers onto the slicing platform of the McIlwain tissue chopper. **Caution:** The tissue chopper blade is extremely sharp. Use caution when placing and adjusting samples.
9. Perform the slicing (500 µm). See Troubleshooting problems 1 and 2.
10. Transfer the Whatman paper with the slices to a new 35-mm petri dish with fresh slicing medium. Carefully remove the human lung organotypic slices using curved tweezers and separate the lung slices. Remove all the Whatman paper from the petri dish.

**Figure 1:**
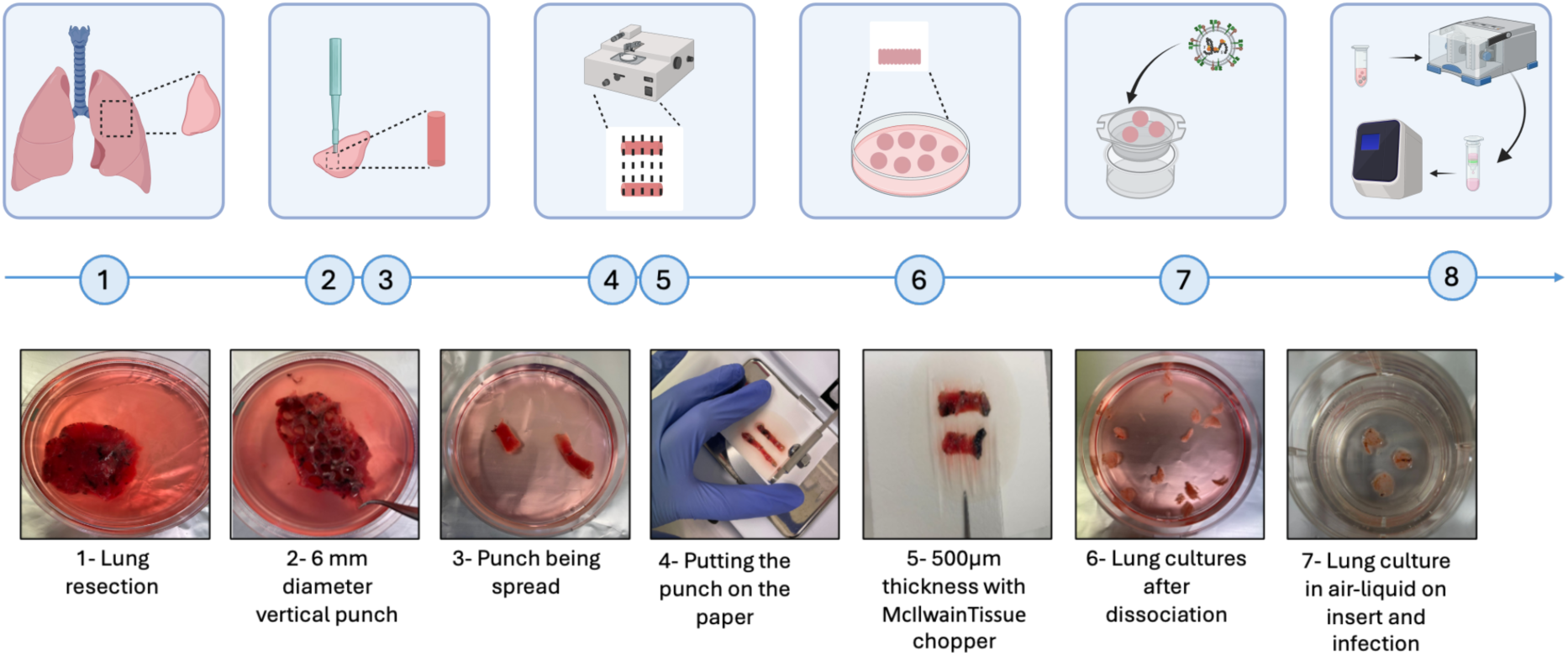
Schematic slicing procedure. Created in BioRender. MATHIEU, C. (2025) https://BioRender.com/ v34z384.

### Transfer onto inserts and culture procedure

#### Timing: 1-3 about 10 min and 20 min for steps 4-7 (variable, depending on the number of slices)

These steps describe how to correctly place tissue cultures on the insert to enable the efficient exchange of nutrients and gas at the air-liquid interface, thereby reducing hypoxia and maintaining tissue viability. The plating procedure is essential to ensure uniform exposure to viral inoculum and/or treatment to preserve reproducible infection/treatment efficiency.

1. Under a safety cabinet carefully remove the medium under the insert using a P1000 Pipetman with filter tips and add 1 mL of fresh medium under the insert before depositing the human organotypic lung slices.
2. Replace the 6 well plate into the incubator under a humidified atmosphere of 5% CO_2_ and 37°C. **Critical step:** This must be completed prior to the beginning of the slicing procedure to avoid drying of the insert membrane and to allow immediate plating of generated slices.
3. Select slices for the culture under the stereomicroscope, slices must be uniform in thickness and round in shape without signs of folding or tearing. Depending on the addressed question, avoid using cultures presenting signs of fibrosis or tar deposits.
4. Carefully place between 2 and 6 slices per insert membrane using the 5 mL truncated pipette mounted on a pipette bulb.
5. Gently remove the excess of slicing medium on the membrane by using a P200 Pipetman avoiding displacement of slices.
6. Incubate the slices at 37°C, under 5% CO_2_ in a humidified atmosphere. See Troubleshooting problems 3 and 4.

### Viability procedure

**Timing:**

**MTT Cell Viability Assay 1-4 about 5 h (including the incubation time).**

**Lactate dehydrogenase (LDH)-Glo Cytotoxicity assay1-4 about 30 min including LDH storage buffer preparation and 1h10 for steps 5-11.**

In this step, we assess tissue viability using complementary methods MTT and LDH, to monitor cellular integrity and metabolic activity. MTT is used as a proxy for tissue viability using quantitative viability assays while LDH quantifies tissue damage by measuring the release of cytoplasmic enzymes into the culture medium.

1. MTT Cell Viability Assay (see Figure 2 a)
  a. Assess the viability of the human lung organotypic slices at 0, 24, 48, and 72 h timepoints.
  b. Transfer huOLC into a 96-well plate and add 70 µL of MTT solution incubate for 4 h at 37°C. Alternatively, prepare a separate plate fully dedicated to viability assay. In this case either immerse slices into MTT solution or directly add 70 µL of MTT solution on the top of each slice to fully cover the huOLC to ensure total exposure of the tissue. **Caution:** MTT is toxic for the skin and eyes, and must be handled in a chemical fume hood and BSL-2/4 conditions are required when working with viruses. Wear gloves and protective eyewear. **Note:** MTT can be prepared in advance, stored at 4°C protected from light and used for 3 weeks.
  c. Transfer each slice into a well of a 48-well plate and cover with 200 µL of DMSO, and triturate to solubilize formazan.
  d. After 10 min incubation at 37°C, transfer 75 µL of DMSO containing formazan, without tissue pieces in a transparent 96-well plate to measure absorbance at 540 nm with a microplate reader (e.g., Tristar 5, Berthold).
2. LDH-Glo Cytotoxicity assay (Figure 2 a) To assess Lactate dehydrogenase (LDH) release in subnatants at 0, 24, and 48 h of culture:
  a. Prepare LDH storage buffer according to [manufacturer instructions](https://www.promega.in/products/cell-health-assays/cell-viability-and-cytotoxicity-assays/ldh-glo-cytotoxicity-assay/?tabset0=1) (see reagents setup section).
  b. Reconstitute LDH with 275 µL of Storage Buffer and mix to make a 1,000 U/mL standard. The LDH standard is aliquoted at –20°C.
  c. Collect 1.5 µL of homogenized subnatant into 148.5 µL of LDH storage buffer to preserve LDH.
  d. Store aliquots at –20°C until day of the assay. The day of LDH quantification:
  e. Thaw samples, LDH detection enzyme mix, LDH storage buffer and LDH standard at room temperature.
  f. Prepare LDH standard dilutions as follows:
    i. Dilute 10 µL of LDH standard in 3.115 mL LDH storage Buffer to reach 3.2 U/mL LDH final concentration.
    ii. Mix 10 µL of 3.2 U/mL LDH standard with 990 µL of LDH storage buffer to reach final concentration of 32 mU/mL.
    iii. Serially dilute 1:2 the 32 mU/mL LDH standard solution by mixing 200 µL of previous concentration with 200 µL of LDH storage buffer until reaching LDH final concentration of 0.5 mU/mL.
  g. Prepare LDH detection reagent by mixing LDH Enzyme Buffer Mix and Reductase: for one reaction, mix 50 µL of LDH assay with 0.25 µL of reductase.
  h. Add 50 µL of sample or standard in a white 96-well plate.
  i. Add 50 µL of detection reagent buffer per well.
  j. Incubate the plate at room temperature for 30 min.
  k. Measure Luminescence using Tristar 5 reader.
3. Cells viability analysis (Figure 2 b)
  a. For each donor pool 2 to 3 slices to ensure having enough cells in total. Note: For count divide the final result by the number of pooled slices.
  b. Incubate each pool for 30 min at 37°C in 500 µL of digestion buffer.
  c. Dissociate cells with approximately 20 aspiration-displacement cycles with a 20G needle.
  d. Filtrate with 40 µm sieve. Wash the sieve three times with DPBS 1X/5% FBS to limit cell loss.
  e. Incubate for 4 min at room temperature with 1 mL of Red Blood Cell Lysis Buffer.
  f. Add 9 mL of DPBS 1X/5% FBS.
  g. Centrifuge 5 min at 500g.
  h. Resuspend in DPBS 1X/5% FBS, 2 mM EDTA.
  i. Centrifuge 96-well plate 5 min at 400g and 4°C.
  j. Meanwhile, dilute the LIVE/DEAD cell reagent mixture 1000 times.
  k. Remove the supernatant by inversion.
  l. Resuspend the cells with 50 µL of LIVE/DEAD cell reagent mixture.
  m. Incubate 25 min at 4°C.
  n. Add 150 µL of DPBS 1X/5%FBS, 2 mM EDTA.
  o. Centrifuge 96-well plate 5 min at 400g and 4°C.
  p. Remove the supernatant by inversion.
  q. Resuspend the cells with 200 µL of DPBS1x/5%FBS, 2 mM EDTA.

The flow cytometry was performed using CytoFLEX cytometer (Beckman Coulter) (without acquisition time).

**Figure 2:**
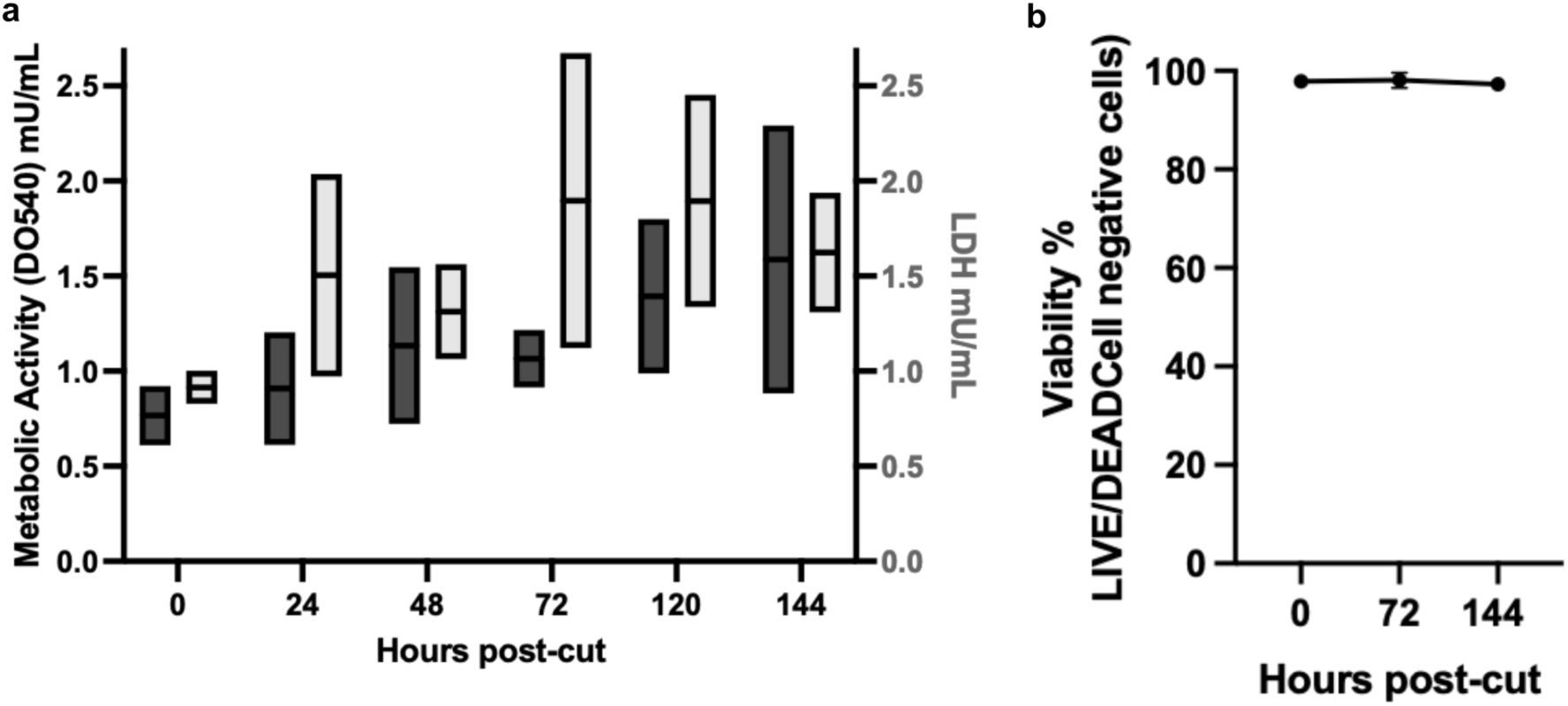
Viability of huOLC (a) Metabolic activity of huOLC was measured by MTT assay at 0, 24, 48, 72, 120 and 144 h post-culture (N=2 separate donors) (in black) and LDH release in subnatants was quantified 0, 24, 48, 72, 120 and 144 h of culture (N=2 separate donors) (in gray). (b) Viability of huOLC were analyzed by LIVE/DEAD Cell labelling at 0, 72, and 144 h of culture (Two separate donors: one sampled at 0 and 72 h, the other at 72 and 144 h).

### Cytoflex count

#### Timing: digestion step (1) 40 min and steps 2-8 about 20 min

This step consists of dissociating the OLCs to recover their constituent cells, which are subsequently quantified by flow cytometry.

Starting from step h) (or step q if LIVE/DEAD is preferred in all staining) described above in “Cells viability analysis” cells were resuspended and analyzed by flow cytometry. Resuspend the cells with 200 µL of DPBS1x/5% FBS, 2 mM EDTA.

1. Perform the flow cytometry using CytoFLEX cytometer(Beckman Coulter) (without acquisition time).
2. Enumerate cells suing flow cytometry-based Precision Count Beads (BioLegend) according to the manufacturer’s instructions.
3. The data were analyzed with Flowjo v.10 (BD Biosciences). Note: only include huOLCs that are viable in the assay.

### Cells phenotype analysis

#### Timing: cells preparation about 8 min (1-3); 25 min of incubation (4); 8 min of wash (5-7) and 8-9 depending on the number of wells

Resuspended cells are stained with fluorophore-conjugated antibodies targeting population-specific markers. Single-cell fluorescence is then analyzed by flow cytometry, allowing identification and quantification of the distinct cellular populations present within the OLCs.

1. After step i) described above in “Cells viability analysis”, cells were stained as described below. The antibodies were listed in table 1. The flow cytometry was performed using CytoFLEX cytometer (Beckman Coulter). Prepare the antibody mixture (the antibodies were listed in table 1).
2. Remove the supernatant of centrifuged plate by inversion.
3. Resuspend the cells with 50 µL of antibody mixture.
4. Incubate 25 min at 4°C.
5. Add 150 µL of DPBS 1X/5% FBS, 2 mM EDTA.
6. Centrifuge 96-well plate 5 min at 400g and 4°C.
7. Remove the supernatant by inversion.
8. Resuspend the cells with 200 µL of DPBS1x/5% FBS, 2 mM EDTA.
9. Perform the flow cytometry in CytoFLEX (Beckman Coulter) (without acquisition time). The data were analyzed with FlowJo v.10 (BD Biosciences). The number of cells was determined using flow cytometry–based Precision Count Beads (BioLegend) according to the manufacturer’s instructions.

**Table 1:**
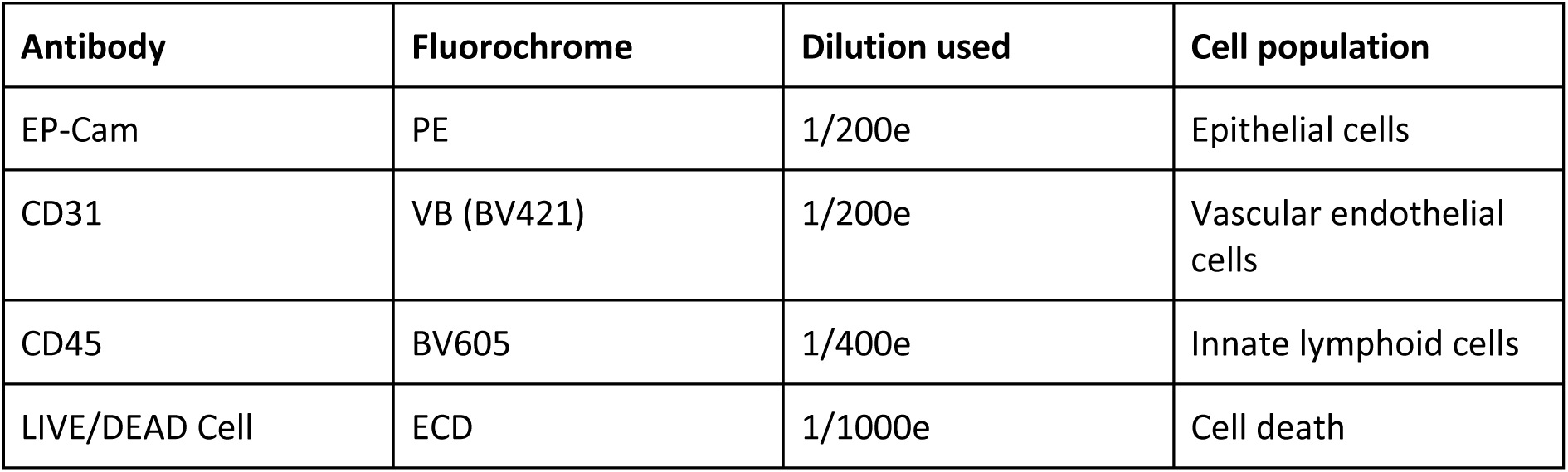
Antibodies and LIVE/DEAD Cell reagent used for cell phenotyping.

| Antibody | Fluorochrome | Dilution used | Cell population |
| --- | --- | --- | --- |
| EP-Cam | PE | 1/200e | Epithelial cells |
| CD31 | VB (BV421) | 1/200e | Vascular endothelial cells |
| CD45 | BV605 | 1/400e | Innate lymphoid cells |
| LIVE/DEAD Cell | ECD | 1/1000e | Cell death |

### Infection procedure

#### Timing: 1-3 about 30 min (depending on the number of slices)

This section describes the viral infection of huOLC by IAV and NiV under controlled conditions to ensure efficient viral entry and preserving tissue viability. Precise parameters are essential for reproducible downstream analyses.

**Note:** All procedures described in the following protocols are performed in biological safety cabinets under sterile working conditions, while wearing personal protective equipment in accordance with BSL-2 and BSL-4 regulations for IAV and NiV, respectively. Refer to your local regulations and permits to determine what biosafety level is appropriate for your strains.

1. Dilute the viral stock in sterile MEM medium for A/Scotland/20/74 (H3N2) or in sterile Opti-MEM medium for rNiV-EGFP to infect the huOLCs. **Caution:** Follow the good laboratory practice and personal protective equipment.
2. Infect the huOLC with 2,000 to 20,000 plaque forming units (pfu) of A/Scotland/20/74 (H3N2) (BSL-2) or 5,000 or 20,000 pfu of rNiV-EGFP (BSL-4). Using low-binding 10 μL pipette tips, dispense a drop of 4 μL of the virus directly on the slice. For the mock control slices, apply the same volume of MEM or Opti-MEM as for the infection. **Critical step:** Ensure that the virus drop is completely and homogeneously distributed on the slice without damaging the tissue.
3. Incubate the infected slices at 37°C under 5% CO_2_ in a humidified atmosphere.
4. For rNiV-EGFP, the fluorescence resulting from viral replication can be followed using a Leica DMIRB microscope, a fluorescence microscope (BSL-4) and Nikon Eclipse Ts2R optical microscope in BSL-2.

### Extraction and RT-qPCR procedure

**Timing:**

**For huOLC infected by NiV: 1-2: 30 min (including 10 min of tissue lyser); 3: 2 h (1 h for the extraction + 30 min rDNAse + 30 min for RNA quantification); 4: 30 min for the preparation of the RT and 26 min for the RT run and 5: 30 min for the preparation of the qPCR and 2 h for the qPCR run.**

**For huOLC infected by IAV: 1-2: 20 min (including tissue lysis using Precellys apparatus); 3: 2 h (1 h for the extraction + 30 min rDNAse + 30 min RNA quantification); 4: 30 min for the preparation of the RT and 1 h 15 min for the RT run; 5: 30 min for the preparation of the qPCR and 1 h 15 for the qPCR run.**

These steps details RNA extraction from infected huOLC and subsequent RT-qPCR analysis to quantify viral and host gene expression. Standardized extraction and RT-qPCR conditions are critical for accurate and reproducible quantification.

Two slices are collected per condition and lysed together to ensure sufficient material for RT-qPCR.

1. Collect the huOLCs at 0, 24, 48, and 72 hours post infection (hpi) and transfer 2 slices per condition into a 2 mL Eppendorf tube containing:
  a. 1 mL of Trizol and 2 stainless beads (5 mm diameter) for rNiV-EGFP
  b. 800 µL of Trizol and 4 stainless beads in Precellys Bertin 2 mL tubes for IAV infection.
2. huOLC lysis program:
  a. For rNiV-EGFP: Use a TissueLyserII (Qiagen) at maximum speed for 10 min to fully dissociate the tissue.
  b. For IAV: Use a Precellys homogenizer with the following protocol performed 3 times: 3 x (15 s at 5196g– 15 s break).
3. Extract total RNA using the Direct-zol RNA Miniprep Kit (Zymo Research), following the [manufacturer’s procedure](https://files.zymoresearch.com/protocols/_r2050_r2051_r2052_r2053_direct-zol_rna_miniprep.pdf). **Critical step:** store purified RNA at –80°C to prevent degradation.
4. Synthetize cDNA with 50 ng of extracted RNA using:
  a. For rNiV-EGFP: the iScript cDNA Synthesis kit (Bio-Rad) according to the [manufacturer’s protocol](https://www.bio-rad.com/sites/default/files/webroot/web/pdf/lsr/literature/4106228.pdf).
  b. For Influenza: the High-Capacity RNA-to-cDNA kit (Thermo Fisher) according to the [manufacturer’s protocol](https://documents.thermofisher.com/TFS-Assets/LSG/manuals/MAN0017977_highcap_cDNA_RT_UG.pdf).
5. cDNA are diluted 1/10. **Critical step:** store the cDNA at –20°C.
6. Perform the qPCR with the Platinum SYBR Green qPCR SuperMix-UDG (Invitrogen) using a StepOne Plus Real-Time PCR System (Applied Biosystems) according to the [manufacturer’s protocol](https://documents.thermofisher.com/TFS-Assets/LSG/manuals/platinum_qpcr_sybr_man.pdf).

**Note:** Use the qPCR primers designed by Beacon Designer (version 8) software or chosen after validation such that their efficacy is close to 100% according to the MIQE checklist^29^. GAPDH (Glyceraldehyde 3-phosphate dehydrogenase) and human Rplp0 are used as housekeeping genes. Run all samples in duplicates and analyze results using StepOne version 2.3 (Applied Biosystems). Normalize all results to the standard deviation (SD) for GAPDH mRNA using GAPDH primers^27^ or Rplp0^9,30^ mRNA and do all calculations using the 2^-ΔΔCT^ model^31^. Perform all the statistical analyses using GraphPad Prism v10.

### Fixation procedures and immunostaining procedure for viral detection

**Timing: 10 min for the fixation (1); step 2-9 about 1 h 30 min; 10 min + overnight for the step 10; 40 min + overnight for 11-12 and 1 h for the last step.**

In this step, we describe fixation and immunostaining procedures used to visualize viral antigens. By staining huOLC after fixation it is possible to determine whether the infection is present or absent.

1. Fix slices infected with rNiV-EGFP with 4% methanol-free paraformaldehyde (PFA) for 14 days. Fix slices infected by A/Scotland/20/74 (H3N2) for 2 h with 4% methanol-free PFA. **Caution:** Carcinogenic, mutagenic or toxic properties. Handle in a fume hood with personal protective equipment.
2. Wash slices 3 times with 1X DPBS, 5 min per wash to remove excess PFA.
3. Gently transfer the slices in a 48-well plate using the 5 mL truncated pipette mounted on a pipette bulb.
4. Saturation of aldehyde groups: add 200 µL/well of 1X DPBS + 0.1% glycine, incubate for 10 min at RT.
5. Repeat step 4. **Caution:** Always check that no bacteria or fungi are present in solutions prior to use.
6. Wash the slices with 1X DPBS.
7. Permeabilize and block non-specific binding sites with 200 µL/well (1X DPBS + 0.5% Triton X-100 + 1% BSA for 2 h or overnight at 4°C).
8. Wash slices in 1X DPBS **Pause point:** Slices can be stored at 4°C.
9. Add the primary antibody, diluted as recommended, in blocking solution and incubate overnight at 4°C with shaking. For IAV-infected huOLCs, viral nucleoproteins were stained overnight at 4°C, using a rabbit anti-Influenza nucleoprotein primary antibody diluted 1:400 in blocking solution.
10. Remove the primary antibody solution and wash 3 times in 1X DPBS for 5 min.
11. Dilute the secondary antibody in blocking solution and incubate slices overnight at 4°C. For IAV-infected huOLCs, the slices were incubated for 2 h at room temperature, with goat anti-rabbit Alexa 488 conjugated secondary antibody diluted 1:1,000 in blocking solution. **Pause point:** The secondary antibody can be left 2 h at room temperature.
12. Wash 3 times: 2 fast washes with 1X DPBS (5 min each) and one long wash (15 min).
13. Incubate huOLCs for nuclei staining for 45 min at room temperature with Hoechst solution diluted 1:1,000 in blocking solution.
14. Mount the slices between slide and coverslip using Fluoromount mounting medium.
15. Observe fluorescence (see Troubleshooting Problem 5).

**Note:** Once mounted, immunostained slices can be stored at 4°C at least for several weeks.

### Titration of IAV infectious particles by TCID_50_ assay

**Timing: 1: 5 min; 2: 35 min (times can vary according to the number of plates); 3-5: 2 days and approximately 1 h 30 to 2 h depending on the number of plates; 6: 2 days; 7-8: 30 min and 9: 10 min**.

This step aims to measure the concentration of infectious viral particles secreted in the subnatants of huOLCs using TCID_50_ titration assay in which serial dilution of huOLC subnatants are used to infect MDCK cells. The resulting TCID₅₀ values determine infectious viral titers and confirm that the huOLCs model is permissive to influenza virus infection and supports effective viral replication in various conditions (time, treatment etc.).

1. Subnatants dedicated to titration are aliquoted and stored at –80°C until the day of titration (see Materials and Equipment section).
2. MDCK cells are seeded at 11,000 cells/well in 96-well plates, in MDCK complete culture medium.
3. Two days after MDCK seeding, the medium is removed and MDCK cells are washed twice with 1X DPBS.
4. Subnatants are diluted in a serial cascade dilution in MEM infection medium (see Materials and Equipment section).
5. 1X DPBS is removed, and cells are inoculated with each dilution (total of 8 replicates for each subnatant dilution).
6. MDCK cells are incubated for 2 days at 37°C, 5% CO_2_.
7. Medium is then removed, and cells are fixed and stained with 0.3% crystal violet + 20% methanol solution for 20 min (see Materials and Equipment section).
8. Crystal violet solution is removed and cells are rinsed three times with 1X DPBS. **Caution:** carcinogenic, mutagenic or toxic properties. Handle in a fume hood with personal protective equipment.
9. Viral titer is determined using the Reed and Muench method^32^.

### Cytokines and chemokines quantification

**Timing: 1: 5 min; 2-3: 5 min; 4: 30-35 min +/− depending on the nature, number and dilution of samples; 5: 1 h 40; 6: 3 h +/− depending on the number of samples for the acquisition on flow cytometer.**

This section describes a protocol to quantify cytokines secretions in huOLC subnatants over time and/ or in various conditions (treatment) using BD Cytometric beads array (CBA) for Human Soluble Proteins Flex Set assay according to the [manufacturer instructions](https://www.bdbiosciences.com/content/dam/bdb/product_assets/product_pdf/kitproduct/pdf_0/558264_558265_Book_Website.pdf).

1. Aliquot and store huOLC subnatants at –80°C. The day of the assay:
2. Humidify the 96-well PALL filter plate with 100 µL of wash buffer.
3. Aspirate wash Buffer gently using a vacuum manifold. **Critical step:** Do not touch the filter with tips. Do not exceed 10’ in Hg of vacuum pressure. Aspirate for 2 to 10 s until wells are drained.
4. Preparing Human Flex Set Standards:
  a. Preparation of the top standard: Mix lyophilized standards from each BD CBA human Soluble Protein Flex Set in a 15 mL falcon tube with 2 mL assay diluent to reach a 5,000 pg/mL dilution.
  b. Incubate top standard solution for 15 min at RT. **Critical step:** do no vortex.
  c. Dilute standard solution by half by mixing 200 µL of standard with 200 µL assay diluent.
  d. Repeat the process until reaching 1:256 dilution, e.g. 8 serial dilution tubes + 1 top standard. **Note:** Additional dilutions can be performed if required.
  e. Prepare a negative control with assay diluent only.
  f. Distribute 40 µL of standard dilution or negative control per well.
  g. Dilute subnatants in Assay diluent to remain within the standard range for each cytokine.
5. Mixing Human Soluble Protein Flex Set Capture beads:
  a. Determine the number of BD CBA human Soluble Protein Flex Sets to be used, as well as the number of samples (including standards) in the experiment.
  b. Vortex each capture beads stock vial for 15 s to resuspend beads.
  c. For each set, and as many as sample, mix 0.8 µL of each cytokine capture bead complete to 40 µL with Capture beads diluent.
  d. Distribute 40 µL of the capture beads mix over 40 µL of sample or standard.
  e. Cover the plate with seal.
  f. Shake the plate for 2 min at 500rpm.
  g. Incubate for 1 h at room temperature, in the dark.
  h. Keep an aliquot of the beads for setup.
6. Preparing Human Soluble Protein Flex Set PE Detection Reagents:
  a. For each set mix and as many as sample, mix 0.8 µL of each Human Soluble Protein Flex Set PE detection reagent complete to 40 µL with detection reagent diluent.
  b. Distribute 40 µL of PE detection mix per well.
  c. Cover the plate with new seal.
  d. Shake the plate for 2 min at 500 rpm.
  e. Incubate for 2 h at room temperature, in the dark.
  f. Aspirate on vacuum manifold until wells are drained.
  g. Add 150 µL per well of wash buffer.
  h. Aspirate on vacuum manifold until wells are drained.
  i. Add 200 µL per well of wash buffer.
  j. Shake the plate for 5 min at 500g to resuspend beads.
  k. Proceed to sample acquisition on flow cytometer (e.g. LSR Fortessa II cytometer, BD Biosciences) according to manufacturer’s instruction (see Troubleshooting Problems 6 and 7).

## Expected outcomes

This protocol enables the generation of standardized, viable, structurally preserved huOLCs that retain the essential architectural and cellular features of the native tissue, making them suitable for a wide range of applications including viral infection studies. A major advantage of this method is the complete elimination of agarose embedding, which significantly reduces preparation time without compromising tissue integrity and preserves resident immune cells such as alveolar macrophages^33^. Furthermore, the use of a tissue chopper ensures a rapid and highly reproducible slicing process, which constitutes a key strength of the technique **(Figure 1)**.

HuOLCs prepared using this protocol are expected to maintain high viability for several days in culture. Tissue viability was assessed by measuring metabolic activity using MTT **(Figure 2 a)** (or MTS assays, not shown), and by monitoring LDH release in the culture medium every 24 h **(Figure 2 a)**. As a complementary approach, cell death was evaluated by LIVE/DEAD staining **(Figure 2 b)** (or TUNEL staining, not illustrated). Together, these assays consistently demonstrate preserved metabolic activity and tissue integrity for up to 6 days (144 h) of culture, validating the suitability of huOLCs as a robust *ex vivo* tissue culture model suitable for infection with human viruses and by extension, with other pathogens such as bacteria or fungi.

As shown in **Figure 3**, the CytoFLEX-adapted protocol allows quantification of cells recovered from the tissue immediately after huOLCs preparation and throughout the culture period.

**Figure 3:**
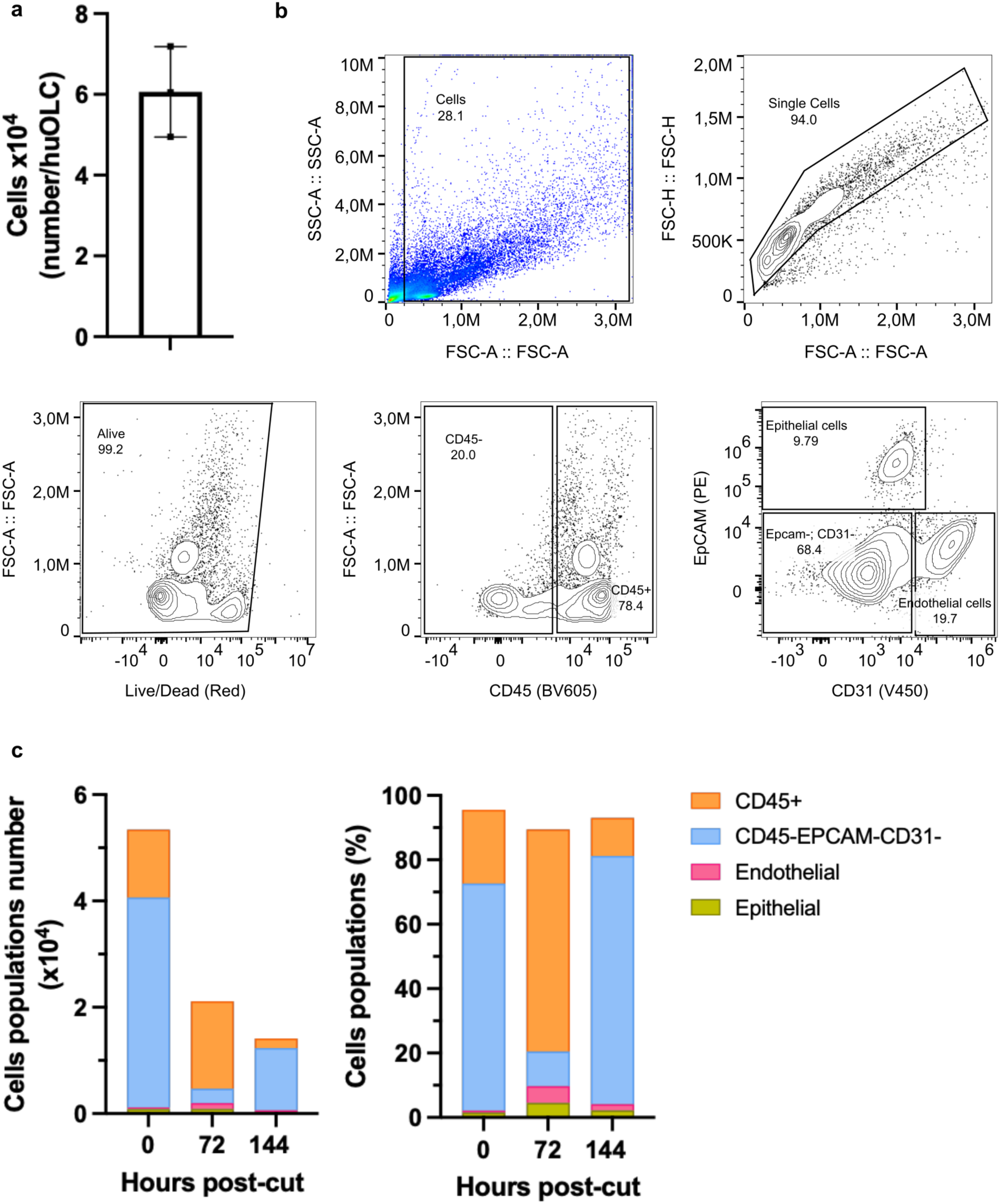
Cells analysis (a) huOLCs were digested for cell dissociation, and cell counting was performed using CytoFLEX at 0 h post-cut. (b) Gating strategy used for the characterization of cell populations in huOLCs. (c) The proportions and numbers of cellular populations present in huOLCs at 0 h, 72 h, and 144 h post-cut were evaluated using CytoFLEX.

One of the primary applications of this model is the infection of human lung slices with viruses spanning biosafety levels from BSL-2 to BSL-4, such as the highly pathogenic Nipah virus (**Figure 4**) or the more commonly encountered IAV H3N2 (**Figure 5**). For example, recombinant Nipah virus expressing EGFP (rNiV-EGFP) **(Figure 4 a)**, can be used to study the early steps of infection by fluorescence microscopy, leveraging the reporter signal to visualize infected cells and monitor viral spread within the tissue. Time-lapse microscopy further allows longitudinal assessment of tissue morphology alongside the progression of EGFP-associated viral infection (**Figure 4 a**). Viral RNA accumulation in tissue cultures can also be quantified by RT-qPCR. HuOLCs are fully permissive to NiV infection, with viral replication reaching a plateau at approximately 48 hpi at the tested inoculum **(Figure 4 b)**. Alternatively, viral growth can be easily assessed by measuring the mean fluorescence intensity from images acquired daily under standardized exposure conditions. This method of analysis confirm the high reproducibility of the infection with rNiV-EGFP between samples, as illustrated at 48h post infection **(Figure 4 c)**.

**Figure 4:**
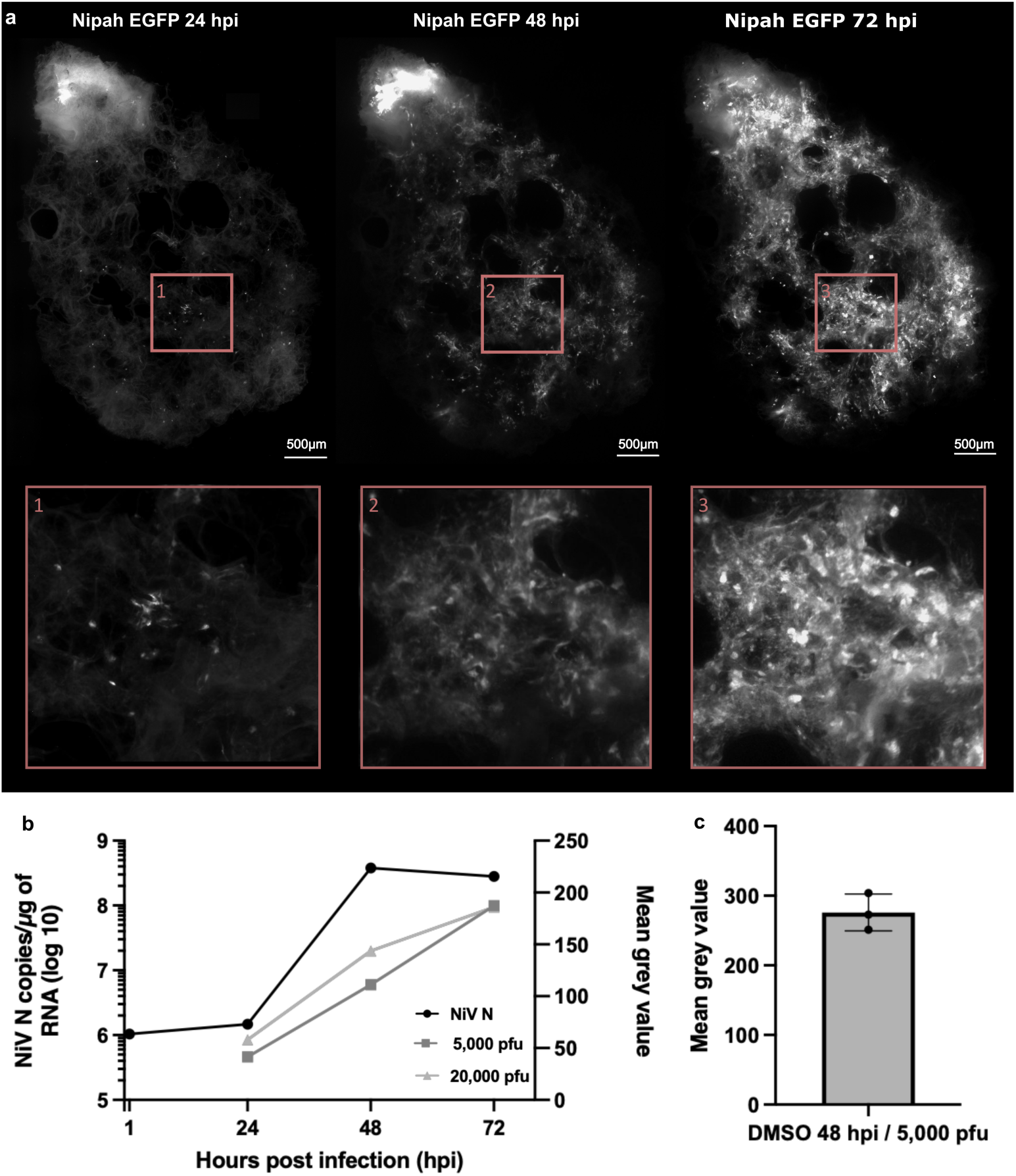
Susceptibility of huOLC to NiV EGFP (a) Representative fluorescence images of huOLC infected with 5,000 pfu of recombinant NiV EGFP. Images were taken at 24, 48 and 72 hpi using a Leica DMIRB microscope at 5x magnification. Exposure times were 2,000 ms for pictures at 24 hpi and 200 ms for pictures at 48/72 hpi. Scale bar = 500 µm. (b) Infection kinetics represented with the NiV replication kinetics in huOLC quantified at 1, 24, 48 and 72 hpi (5,000 PFU of rNiV-EGFP). huOLC were lysed and total RNA extracted, and viral replication was quantified by RT-qPCR targeting the NiV nucleoprotein (N) gene. In parallel, EGFP expression was determined by fluorescence microscopy and results show the mean grey value of huOLC infected with 5,000 PFU or 20,000 PFU at 24/48/72 hpi. Threshold setting: 90-65565, mean grey value obtained on a specific ROI. (c) Mean grey value of huOLC infected with 5,000 PFU NiV EGFP at 48 hpi. Threshold setting: 90-65565, mean grey value obtained on a specific ROI. Data represent mean N=3 +/−SD.

**Figure 5:**
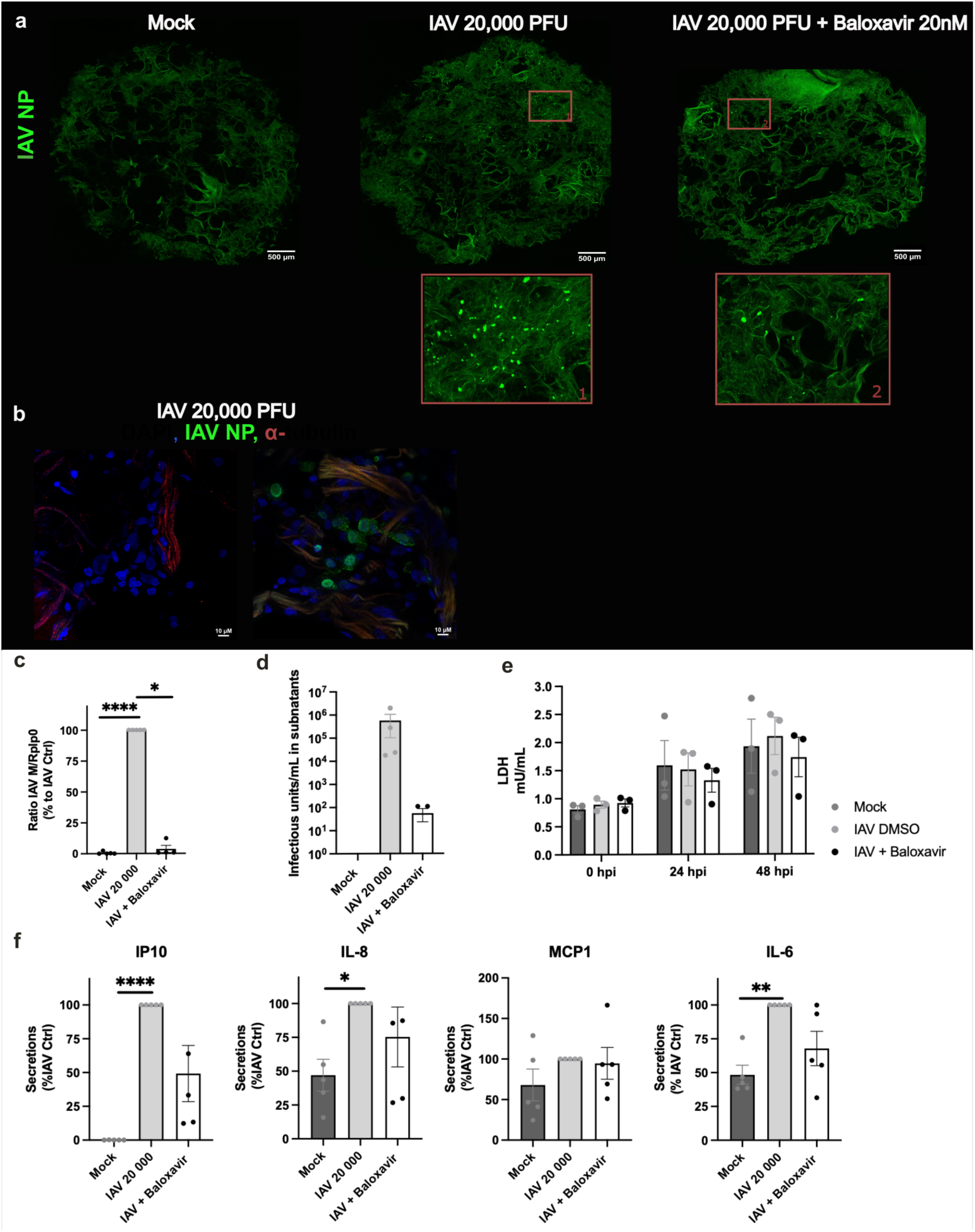
Susceptibility of huOLC to IAV H3N2/Scotland strain. (a) Representative immunofluorescence images showing IAV nucleoprotein expression at 48 hpi, in huOLCs that were either not infected, infected with 20,000 pfu of A/Scotland/20/74 (H3N2) virus, or infected and treated with 20 nM Baloxavir. Images were taken using a spinning Disk confocal microscope CQ1 Yokogawa CQ1 confocal microscope, at x10 magnification. Exposure times were 0.5 s. Scale bar = 500 nm. (b) Confocal microscopy analysis of IAV-infected huOLC at 48 hpi following infection with 20,000 pfu of Influenza A/Scotland/20/74 (H3N2) virus. Viral nucleoprotein (green), nuclei (blue) and α-tubulin (red) are shown. huOLC were stained overnight at 4°C with anti-Tubulin (1/500) and anti-NP-FITC (1/50) antibodies. The anti-mouse-Alexa Fluor 647 secondary antibody was applied for 2 h at room temperature for α-tubulin staining. Images were acquired using a Leica SP8 confocal microscope at x63 magnification, exposure time 1 to 2 min and 3D reconstructions were generated with Leica LasX Life Sciences Software. Scale bar = 10 µm. (c) IAV replication in huOLCs at 48 hpi was quantified by RT-qPCR targeting the viral Matrix (M) segment and normalized to the housekeeping gene Rplp0 gene following tissue lysis and RNA extraction. (d) Quantification of infectious viral particles in huOLC culture at 48 hpi following infection with 20,000 pfu of A/Scotland (H3N2) strain, in the presence or absence of 100 nM of Baloxavir. Viral production was determined by TCID50 titration on MDCK cells. (e) Measurement of huOLCs viability by quantification of lactate dehydrogenase (LDH) secretion into the culture medium, as a proxy of cell death. Supernatants were collected at 0, 24, and 48 hpi. (f) Quantification of cytokine and chemokine secretion in the basolateral compartment at 48 hpi. HuOLCs were infected with 20,000 pfu of IAV A/Scotland/20/74 (H3N2) strain and treated or not with 100 nM Baloxavir. Cytokines and chemokines were quantified using bead-based immunoassays (CBA).

The susceptibility of huOLCs to IAV infection was characterized at 48 hpi using complementary approaches: immunofluorescent detection of viral nucleoprotein, quantification of viral genome replication by RT-qPCR, and titration of infectious particles in culture supernatants by TCID_50_ assays **(Figure 5 a-d)**. As observed in both NiV– and IAV-infected cultures, a degree of intrinsic tissue autofluorescence may be detected during imaging mainly when working with green fluorescence.

Beyond infection studies, huOLCs can be used to evaluate the antiviral activity of diverse therapeutic molecules. For instance, Baloxavir, a well-characterized inhibitor of IAV polymerase, exhibited significant antiviral efficacy at the concentrations of 20 and 100 nM **(Figure 5 a-d)**. Treatment of IAV-infected huOLCs with Baloxavir did not affect tissue viability as indicated by stable LDH secretion over time **(Figure 5 e)**. In parallel, inflammatory cytokines (IL-6 and IL-8) and chemokines (IP10 and MCP1) secreted into the basolateral compartment can be quantified over time by ELISA (**Figure 5 f**). This allows not only characterizing the inflammatory tissue response to infection but also evaluating the anti-inflammatory efficacy of the tested treatments. It is important to note that for milder infections, depending on the strain of IAV or SARS-CoV-2, substantial donor-to-donor variability in both cytokine and viral quantification can be observed. Baseline cytokine levels vary widely across donors and inevitably influence the magnitude and kinetics of responses measured following infection. Similarly, infection efficiency can differ markedly between donors, likely reflecting individual’s history (e.g. prior infections, vaccination and environmental exposures, *etc.*) compared to the relatively uniform responses observed in syngeneic mouse models. This effect can be particularly pronounced for Influenza virus, for which past infections and vaccinations are suspected to significantly shape immune imprinting and, consequently, the responsiveness of organotypic lung cultures. Although this inter-individual variability increases experimental complexity and may challenge reproducibility, it also represents a major strength of the model. By capturing the inherent heterogeneity of human infectious history –something largely absent from traditional experimental systems-these human organotypic cultures provide a physiologically relevant platform for evaluating novel therapies and assessing their robustness across diverse human backgrounds.

## Quantification and statistical analysis (optional)

All the quantifications and statistical analyses were performed using GraphPad Prism, FlowJo and ImageJ. Details of the number of biological replicas (n) and statistical tests used are indicated in the corresponding figure legends.

Flow cytometry data were analyzed using FlowJo. Cell populations were identified by sequential gating (Figure 3).

Quantification of fluorescence signal for NiV was performed by measuring the mean gray value within manually defined and applied to all images within the same experiment. Background signal was subtracted prior to quantification. When images were acquired using different exposure times, signal intensities were normalized by applying a correction factor corresponding to the exposure time ratio using Process > Math > Divide in ImageJ (*).

Fluorescence images were processed and analyzed using ImageJ/Fiji. When necessary, tiled images were reconstructed using the stitching plugin.

(*) DMSO condition – 48 hpi (5,000 pfu): Images were acquired with an exposure time of 300 ms, whereas comparable conditions were acquired at 200 ms. To normalize signal intensities across images, a correction for exposure time was applied. Signal intensities were normalized by dividing the measured intensity values by the exposure time ratio (300 ms / 200 ms = 1.5) using Process > Math > Divide in Image J.

Kinetics – 5,000 pfu: For kinetics analyses, images acquired at 24 hpi were obtained with an exposure time of 2,000 ms while at 48 hpi and 72 hpi, images were acquired with an exposure time of 20 ms. Due to these differences, a correction was applied to the 24 hpi images to allow comparison across time points. Signal intensities at 24 hpi were divided by 10 using Process > Math > Divide in Image J.

Kinetics – 20,000 pfu: Similarly, for 20,000 PFU condition, images at 24 hpi were obtained with an exposure time of 2,000 ms whereas images at 48 hpi and 72 hpi, images were acquired at 20 ms. A correction was therefore applied to the 24 hpi images by dividing signal intensities by 10 using Process > Math > Divide in Image J to normalize signal value and enable direct comparison between time points.

## Limitations

A limitation of this procedure is that the lung resections used came from different anatomical areas of the lung which may contain varying proportions of cells, and so influence infection, as well as altered regions, sometimes fibroses or pigmented area such as “black aggregates” linked to exposure to tobacco/pollution/tar. This tissue heterogeneity may influence the permissiveness of huOLC to infection. It is important to have inclusion and exclusion criteria that will allow a better appreciation of the number of samples to include for statistical analysis.

## Troubleshooting

### Problem 1

Core tissue sample can stick to the chopper blade of the tissue chopper due to the tissue dryness (Step 1).

### Potential solution 1

Moisten the “core tissue sample” and the Whatman paper.

### Problem 2

If the blade does not cut the Whatman paper then it will not slice deeply enough the tissue (Step 1).

### Potential solution 2

It is important to perform a test with the blade before slicing the “core tissue sample”, using only Whatman papers and medium. If the blade does not cut enough, you must change the blade. This phenomenon may also be visible after a few uses of the blade, indicating that you need to change the blade.

### Problem 3

Sometimes, it is possible to see contamination on huOLCs in the form of bacteria and fungi. This contamination can be visible to the naked eye as well as under a microscope. This contamination can be due to the medium but also the air from dissection environment (Step 1).

### Potential solution 3

To avoid this problem, it might be advisable to perform the dissection under a PSM, filter the different media used, clean the room/instruments, in particular the tissue chopper/biopsy punch/bulb, and turn off the ventilation/air conditioning in the dissection room.

### Problem 4

Sometimes huOLCs are not homogeneous, due to heterogenous “core tissue sample” extraction, which can cause problems during infection and analysis (Step 2).

### Potential solution

It is important to choose “core tissue samples” carefully in order to obtain homogeneous huOLCs. This selection must be done under a stereomicroscope to minimize variability.

### Problem 5

The strong natural autofluorescence of lung tissue can overlap with the antibody conjugated fluorophores emitting between ∼450 and 550 nm.

### Potential solution

Select a fluorophore that is distant to lung natural autofluorescence wavelength. Use a confocal microscope

### Problem 6

Cytokines and/ or chemokines are not detected into huOLC subnatants. This can be due to sample being too diluted.

### Potential solution

Try various sample dilutions until finding the optimal one.

### Problem 7

Cytokine level in samples is above top standard value. The sample is not diluted enough.

### Potential solution

Try various dilutions of the sample until finding the optimal set up.

**Note:** Sample dilution can vary depending on the cytokine quantified.

## Resource availability

### Lead contact

Further information and requests for resources and reagents should be directed to and will be fulfilled by the lead contact, Cyrille Mathieu.

### Technical contact

Technical questions on executing this protocol should be directed to and will be answered by the technical contacts, Lola Canus, Florentine Jacolin or Virginie Vasseur.

### Material availability

This study did not generate new unique reagents.

### Data and code availability

No new data/code was generated in this study. Any additional information required to reanalyze the data reported in this paper is available from the lead or technical contact upon request.

## Acknowledgments

The project was supported by ANR-ASTRID (ViroMetaBlock project (ANR- 22-ASTR-002) funded by the Agence Nationale de Recherche and the Agence Innovation Défense (AID) as well as by ISIDORe. This work also received support from the French government through the France 2030 program and the ANRS | MIE, specific project ANRS-23-PEPR-MIE 0008 (NIPAH-LISA). LC was supported by a Ph.D fellowship from INSERM, the Direction Générale de l’Armement (DGA), and the Agence Innovation Défense (AID). FJ was supported by a Ph.D. fellowship from the Agence Nationale de Recherche (Program Fluccinate; ANR-22-CE15-0001-01). The salary of EO was funded by ANRS | MIE, specific project ANRS-23-PEPR-MIE 0008 (NIPAH-LISA).

The ISIDORe research program is co-funded by the European Union and has received funding from the European Union’s Horizon Europe research and innovation program under grant agreement number 101046133. Views and opinions expressed are those of the author(s) only and do not necessarily reflect those of the European Union or the European Research Executive Agency (REA). Neither the European Union nor the granting authority can be held responsible for them.

We are grateful to the CHRU of Tours and HCL of Lyon for providing access to human lung resection samples, and to the patients for their informed consents. We also acknowledge the P4 Jean Mérieux laboratory for its essential support in biosafety, biosecurity and high-containment experimentation. We thank Dr Alexandre Lalande and Clémence Jacquemin for their valuable assistance throughout this study. We thank the imagery platform (LYMIC-PLATIM) of SFR Biosciences Gerland-Lyon Sud and especially Elodie Chatre for her help. We also thank the Plate-Forme IBiSA des Microscopies, PPFASB, Université de Tours and CHRU de Tours. All statistical analyses were performed with GaphPad v10.

## Author contributions

Conceptualization: CM, MST, FA, LPC, POV. Methodology: LC, FJ, VV, CM. Software: LC, AB, VV, AC, CM Validation: MS, AG, AL, CD, DS, LPC, POV, LC, FJ, VV, AC, FA, AE, CM. Formal analysis: LC, FJ, VV, AC, EO, AB, AAG, CM. Investigation: LC, FJ, VV, AAG, AC, AG, MST, LPC, POV, CM. Resources: MST, AG, AL, AE, FA, VL. Data curation: LC, FJ, VV, AAG, AC, AG, MST, LPC, POV, CM. Writing – original draft: LC, FJ, VV, AC, AB, LPC, MST, AG, POV, CM. Writing – review & editing: LC, FJ, VV, AC, AB, LPC, MST, AG, AL, CD, DS, VL, POV, CM. Supervision: CM, MST, LPC, POV. Project administration: MST, LPC, AG, POV, VL, CM. Funding acquisition: CM, MST, LPC, POV, VL. All authors have read, commented and approved the final manuscript.

